# Twister3: a simple and fast microwire twister

**DOI:** 10.1101/727644

**Authors:** Jonathan P. Newman, Jakob Voigts, Maxim Borius, Mattias Karlsson, Mark T. Harnett, Matthew A. Wilson

## Abstract

We present Twister3, a microwire twisting machine. This device greatly increases the speed and repeatability of constructing twisted microwire neural probes (e.g. stereotrodes and tetrodes) compared to existing options. It is cheap, well documented, and all associated designs and source code are open-source. Twister3 is of interest to any lab performing twisted microwire neural recordings, for example, using tetrode drives.

## 1 Introduction

Since their introduction [1], twisted wire probes (TWPs; e.g stereotrodes [1] and tetrodes [2, 3]) have been a reliable method for obtaining single-unit extracellular spiking data in freely moving animals. They are cheap (~5–10 USD/m using Sandvik PX000004), small enough to cause minimal inflammation, and sufficiently biocompatible to be used over many months [4, 5]. Their contacts are close (~10–20 *μ*m) and therefore allow much improved unit separability compared to single wire probes [6]. They are mechanically flexible, such that they move with neural tissue, rather than behaving like a rigid, skull-coupled beam. This mitigates “drift” [7, 8] and improves long term stability compared to with low-density silicon probes^1^, allowing units to be tracked continuously over multi-day timescales [4]. Although the introduction of modern silicon [9, 10] and carbon-fiber probes [11] offer major advances in terms of channel density and size, respectively, and rival microwires in terms of mechanical flexibility [10], TWPs will remain a ubiquitous recording method for the foreseeable future due to their simplicity, good performance, and low cost.

Although simple to make [12], constructing TWPs is a tedious and time-consuming process. For modern, easy-to-assemble microdrive designs [5], making TWPs is a rate-limiting step. Typically, TWPs are made in three actions [12]:

1. **Folding**: insulated tungsten or nickel-chrome (‘nichrome’) resistance wire is drawn from a spool and folded by hand 1 or 2 times to make stereo- or tetrodes, respectively.
2. **Twisting**: the folded wire is draped over a smooth metal rod. The free end of the wire bundle is loosely coupled to a motor armature using a mechanical^2^ or magnetic^3^ mechanism. The motor then twists the wire bundle into a helix.
3. **Fusing**: A hot air gun is used to fuse the insulation on the wire helix, forming a springy, implantable probe.

Recently, SpikeGadgets^4^ introduced a tetrode twisting machine that provides an arrangement of four pre-wound wire bobbins that allows the user to draw a multi-wire bundle without the folding step^5^. Because folding is the most time consuming part of the TWP making process, this device greatly improves the speed at which TWPs can be created. Additionally, this method minimizes human wire handling, which is beneficial because wire ends up in direct contact with neural tissue and is difficult to clean. Despite these advantages, the high-price of its custom-designed components (~10,000 USD) and complexity of this device has hampered its adoption. Inspired by the SpikeGadgets design, we collaborated to create an open-source twisting machine that is approximately ten times cheaper and ten times faster (Fig. 1 (A)). Our device uses a high-speed stepping motor and modern micro-stepping driver to increase twisting speed while maintaining precise control of motor acceleration and smooth motor actuation. A new counter-balanced, auto-aligning spring system provides constant tension on tetrode wire during twisting. Our design allows a moderately trained (~1 hr of experience with device) operator to make ~70 TWPs per hour.

**Figure 1:**
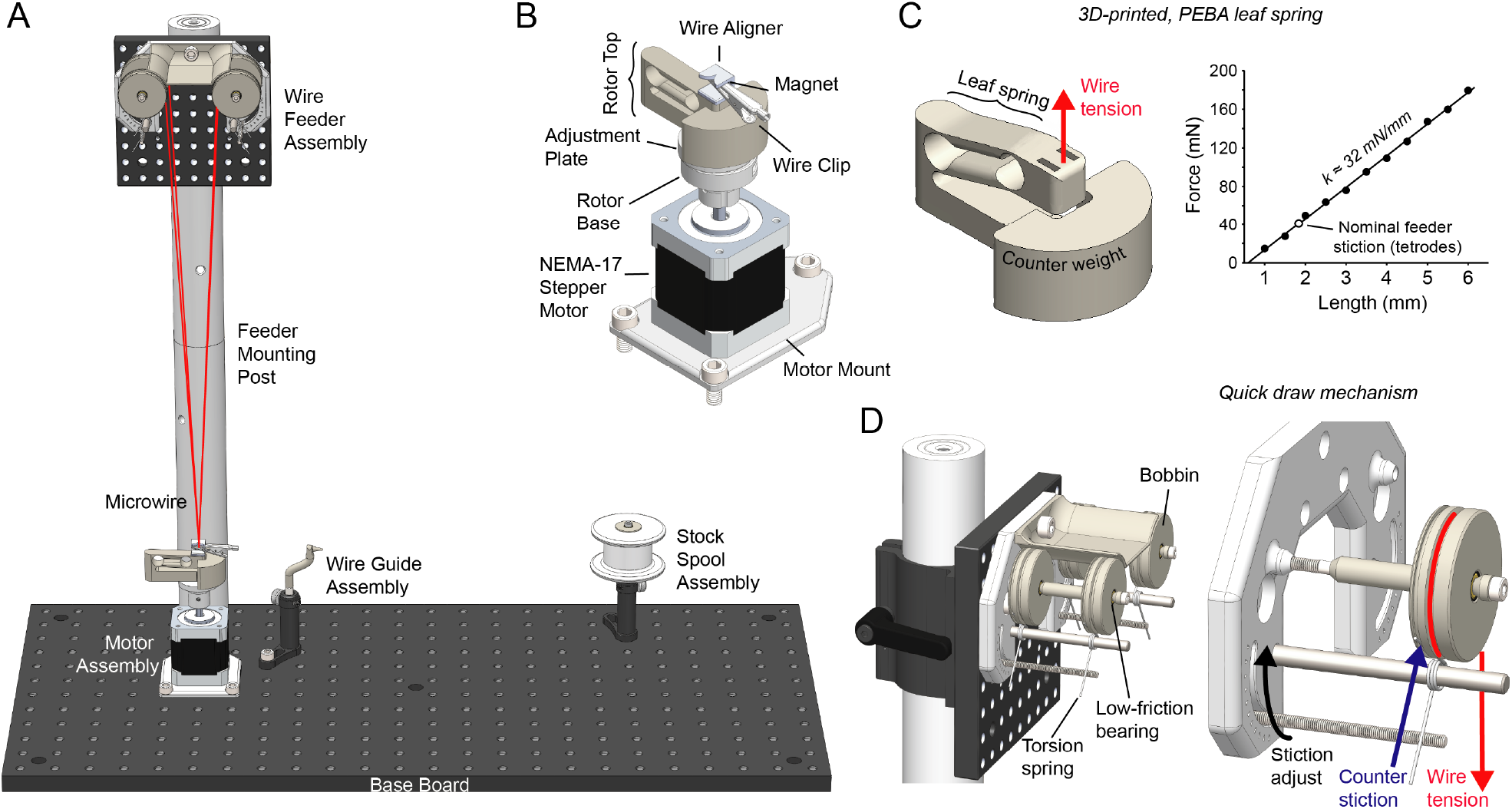
**(A)** Overview of Twister3’s mechanical components. The motor and wire feeder assemblies are used to rapidly construct TWPs by drawing a wire bundle from the feeder, clipping it to the motor, and performing a twist. Additionally, the motor, wire guide, and stock spool assemblies are used to load wire onto the bobbins in the feeder after they are depleted. **(B)** Motor assembly. A NEMA-17 stepper motor is used to twist TWPs and reload bobbins. The wire clip, alignment jig, and magnet allow the wire bundle to be rapidly and reliably linked to the motor. The rotor base and adjustment plate allow one-time adjustment to achieve perfect alignment between the wire bundle and the motor axis. **(C)** *(left)* The 3D printed leaf spring showing deformation under tension. The shape of the spring permits approximately vertical deformation of the wire attachment point so that the center axis is maintained as the bundle is shorted due to twisting. *(right)* Spring tension as a function of vertical deformation. Best fit line indicates a spring constant of 32 nM/mm. The white dot indicates the spring deformation needed to oppose the wire-feeder’s stiction setting for TWPs made in our lab. **(D)** *(left)* Wire quick draw mechanism. *(right)* Isolated single bobbin indicating the wire tension, due to the leaf spring in (C), and counter stiction due to the adjustable torsional spring.

Here we present descriptions of how this device works, materials and assembly information, and electronics designs. We provide detailed usage instructions, an exploration of probe mechanics with respect to twisting parameters, TWP construction time measurements, and show data obtained with tetrodes made using this machine. All designs and source code associated with this project can be found on the Twister3 git repository^6^.

## 2 Notable Design Elements

### 2.1 3D-printed Leaf Spring for Fast Twisting

In order to achieve straight TWPs, wire must be twisted together while under tension. In all other existing designs, a weight is hung from the wire bundle and a motor is loosely coupled to the weight in a way that does not constrain axial motion. This provides constant tension on the wire bundle due to gravity while allowing the helix to decrease in length as it is twisted. A variant of this method uses magnets hold the weight in place^7^, but, in our experience, this is unnecessary and is prone to causing wire breakage due to the nonlinear force/distance relationship of magnetic attraction.

A key design criterion for our device is that TWPs need to be turned very quickly. We aimed for <10 seconds of twisting time (time when motor is in motion) per TWP. Assuming 100 total revolutions (upper estimate for tetrodes) this translates to an average turn rate of 600 RPM. Because previous methods rely on loose motor coupling, they were unsuitable to meet our speed requirements. The high centripetal forces involved in rapid turning inevitably leads to instability of the coupling mechanism, causing the bundle to vibrate wildly. Therefore, we sought to rigidly constrain the motor and wire bundle in the turning plane, but still maintain freedom in the axial direction to allow bundle tensioning and shortening during the turning process. To meet this goal, we made use of selective laser-sintered polyether block amide (PEBA) to create a monolithic, combined leaf spring/wire-retention mechanism (Fig. 1 (B)). PEBA provides rubber-like mechanical qualities resulting in a spring-constant low enough for use with tetrode wire (~32 mN/mm in the relevant range of motion (Fig. 1 (C)).

### 2.2 Auto-aligning Wire/Motor Interface

To interface the microwire bundle with the motor, the bundle is clipped together using a standard, ferrous alligator clip that has been coated in insulating shrink wrap. The bundle is then drawn towards the motor while the leaf spring is lifted with with the user’s free hand. While the leaf spring is under tension, the alligator clip is attached to a strong neodymium magnet on the leaf spring assembly, which provides the bundle-to-motor linkage (Fig. 1 (A, B)). During this process, a 3D-printed alignment jig on top of the magnet automatically guides the microwire bundle in line with the motor’s axis of rotation. The leaf spring is then slowly lowered until it reaches equilibrium with upward wire tension. At this point the motor can be turned.

Because of print tolerances and the fact that the leaf spring does not deform exactly vertically, the resting position of wire bundle will likely not be perfectly co-linear with the motor axle. To account for this, the motor mount consists of a two pieces: the base, which is rigidly concentric with the motor shaft, and an alignment plate. The alignment plate is friction fitted within the base and provides ~2 mm of omnidirectional planar adjustment (Fig. 1 (B)). This plate should be moved until the resting position of the wire bundle is in line with the motor axis.

This wire attachment mechanism has two advantages over existing designs. First it can be used rapidly because of magnetic coupling. Second, because the microwire bundle is automatically aligned with the motor’s axis of rotation, high-turn rates do not result in oscillations or instability. We have found that this is a critical feature in order to produce straight, even-pitch, and consistent TWPs using very fast turn speeds.

### 2.3 Quick-draw Wire Feeder

To further increase the speed at which TWPs can be made, microwire needs to be drawn and attached to the motor quickly. Ideally, this process should occur with as few separate actions being taken by the operator as possible. To facilitate the rapid draw of wire from stock spools, we designed a torsion spring-based feeding assembly that allows wire to be quickly drawn from stock feeder bobbins (Fig. 1 (C)). This mechanism applies enough friction to feeder bobbins to counter increased wire tension during a twist, transferring all slack compensation to the leaf spring (Fig. 1 (B)). The holding (stiction) force of this mechanism is adjustable to account for the elastic deformation of different wire materials. We have found that, when working with standard tetrode wire^8^, the 2nd stiction setting (corresponding to a threshold of ~11.5 N per spool), is adequate to counter wire tension and therefore prevent spools from improperly feeding during a twist. However, the proper stiction setting is dependent on the microwire material and will need to be adjusted for other wire types.

### 2.4 Motor Control Hardware for Smooth Turning

To obtain precise control over motor acceleration, speed, and position we to used a bipolar stepper motor to perform wire twisting. We have found that, due to their discretized motion, stepper motors can vibrate resonantly with taut microwire, resulting in irregular twists and wire damage. To overcome this issue, we drive our motor using an advanced microstepping driver (Fig. 2 (A)); Trinamic TMC2130). In our case, we use 200 steps/revolution (1.8°) motor. Microstep commands from the microcontroller are provided at 16 microsteps/step, which are further interpolated to 256 microsteps/step by internal driver circuitry. This results in a motor update resolution of 3,200 microsteps/revolution (0.113°), and a motion discretization of 51,200 steps/revolution (0.007°). Therefore, the motor operates approximately as smoothly as a continuous DC motor but with much improved motion control dynamics.

**Figure 2:**
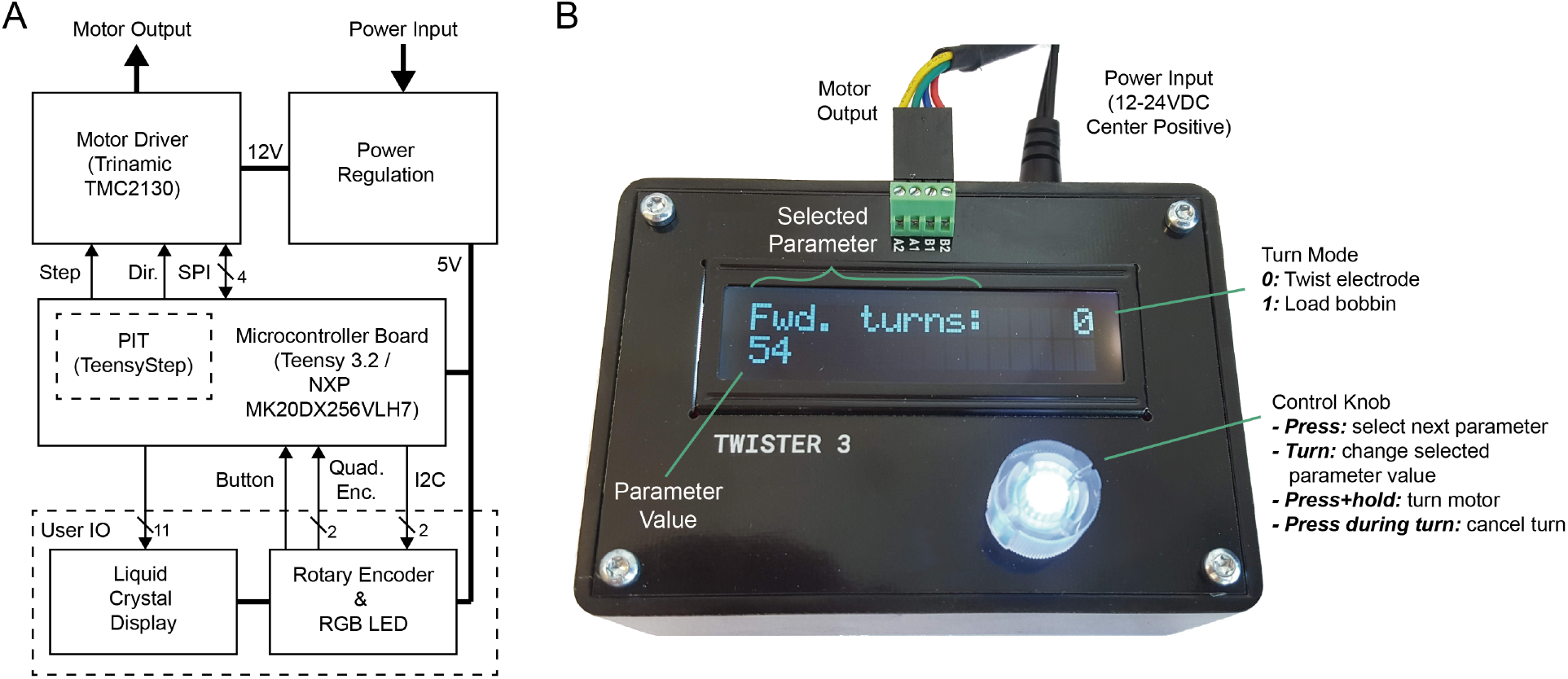
**(A)** Control electronics block diagram. All user IO is provided via a combined rotary encoder and button. The Teensy’s NXP MK20DX256VLH7 microcontroller provides a programmable interrupt timer (PIT) to control step commands to the motor driver, independent of nominal operation. **(B)** Control box with callouts showing features, connections, and controls.

We used an Arduino-compatible Teensy 3.2^9^ microcontroller module to perform step timing calculation. Its NXP MK20DX256VLH7 Cortex-M4 includes an integrated programmable interrupt timer (PIT) which is used to provide jitter-free step commands to the motor driver while acceleration calculations are performed^10^.

## 3 Usage

The following sections provided detailed instructions for using Twister3 to load wire and make TWPs. This content is aided by an instructional video available on YouTube (Appendix A).

### 3.1 Using the Control Box

The control box (Fig. 2 (B)) is powered using a 12V DC center-positive barrel jack that supplies at least 1.5A. It has a single user input: a control knob consisting of a combined quadrature rotary encoder and tactile push-button. This knob permits the following user actions:

- **Press:** cycle through different settings (forward turns, backward turns, turn speed, turning mode)
- **Turn:** increment or decrement the selected setting depending on turn direction.
- **Press and hold:** execute the turn sequence using the current settings
- **Press during motion:** cancel the twist and stop the motor immediately.

The control box is used to perform two tasks: twisting electrodes (turn mode 0) and loading bobbins with microwire (turn mode 1). The turn mode is selected and changed using the dial on the controller. The selected turn mode is shown in the upper right corner of the liquid crystal display (LCD). After a mode is selected, all turning parameters (speed, forward and backwards turns) pertain to that mode only. All parameters are stored in non-volatile memory when a turn is started by pressing and holding the control knob. In the following sections, we detail how to use the mechanical components for making TWPs and loading bobbins with stock wire.

### 3.2 Loading Bobbins

Before twisting electrodes, the bobbins on the wire feeder assembly must be loaded with microwire (Fig. 3A). The following steps detail the bobbin loading procedure:

1. Remove the wire shield by removing its M6-retention screw.
2. Remove one set of bobbins by unscrewing the M3 bolt that serves as the axle.
3. Take the bobbins and spacers off the axle.
4. Remove any remaining microwire from each bobbin and ensure they are clean of dirt and debris.
5. Remove the leaf spring to the base rotor.
6. Place a bobbin on the motor using the embedded magnets (Fig. 3 (A1))
7. Adjust the position of the wire guide such that the tip points directly into the center of the wire groove on the bobbin. The tip of the wire guide should be a few millimeters away from this groove (Fig. 3 (A1)).
8. Feed the tetrode wire from the stock spool through the wire guide and wrap once around the bobbin in its center groove (Fig. 3 (A2)).
9. Set the controller to “mode 1”.
10. Select the desired loading speed. We have found that 100 RPM works well for our wire.
11. Select the desired number of turns to load the bobbin. The circumference of the bobbin is ~10 cm. The length of wire loaded on the bobbin is therefore *turns* × 10 *cm*. Including wastage, this results in a conservative estimate of 1 TWP per turn (Table 1).
12. Start the turn and wait until it is finished. *Be careful not touch moving parts during this process*: the microwire needs to have constant tension to ensure it is properly loaded on the bobbin.
13. Repeat the process for the remaining bobbins.
14. Put the bobbins back on their axle on the wire feeder assembly. Loose wire ends should point inward on both sides of the assembly.
15. Replace the wire shield.

**Figure 3:**
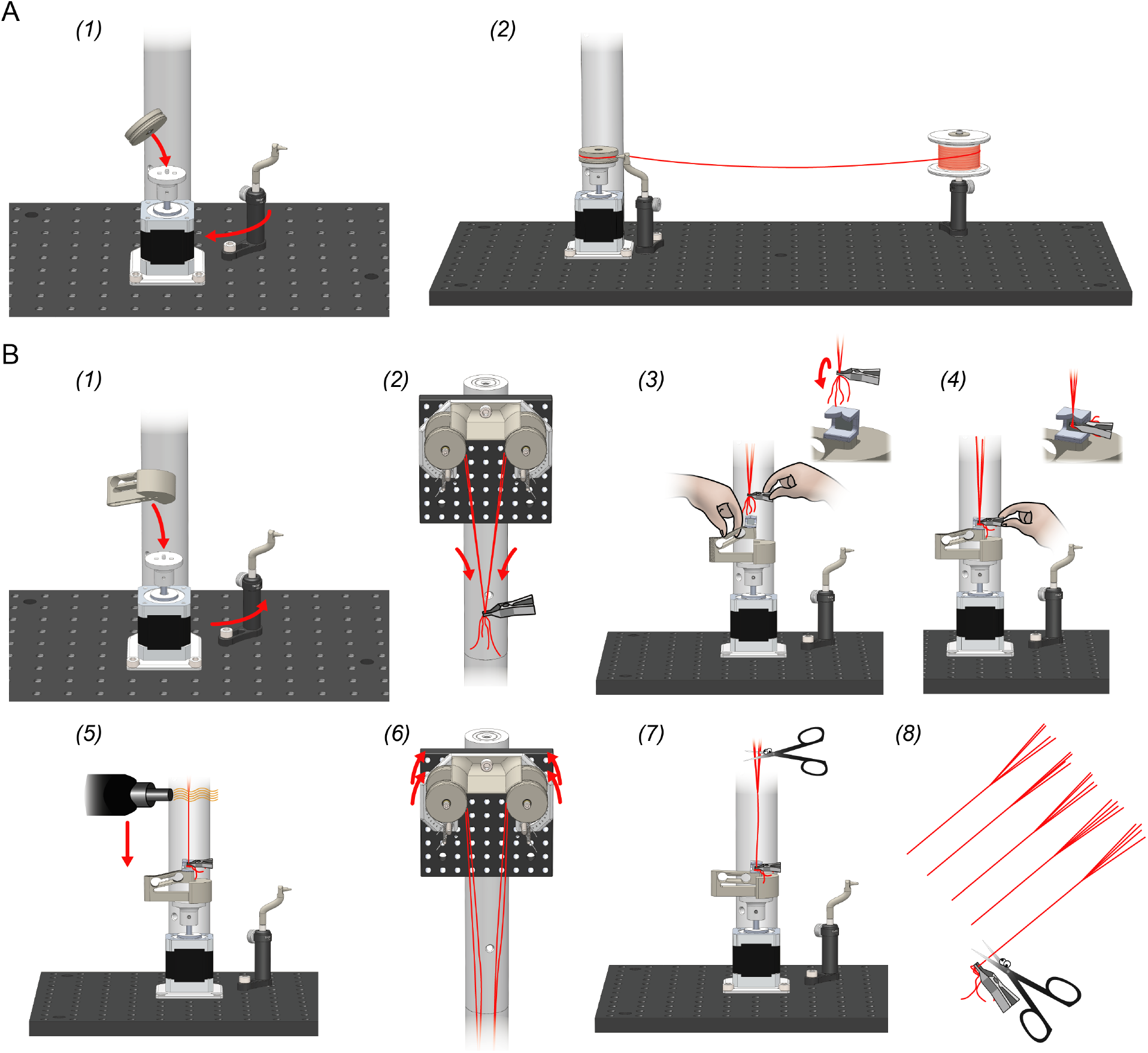
**(A)** Bobbin loading procedure. **(B)** TWP construction procedure. See text for details.

**Table 1:**
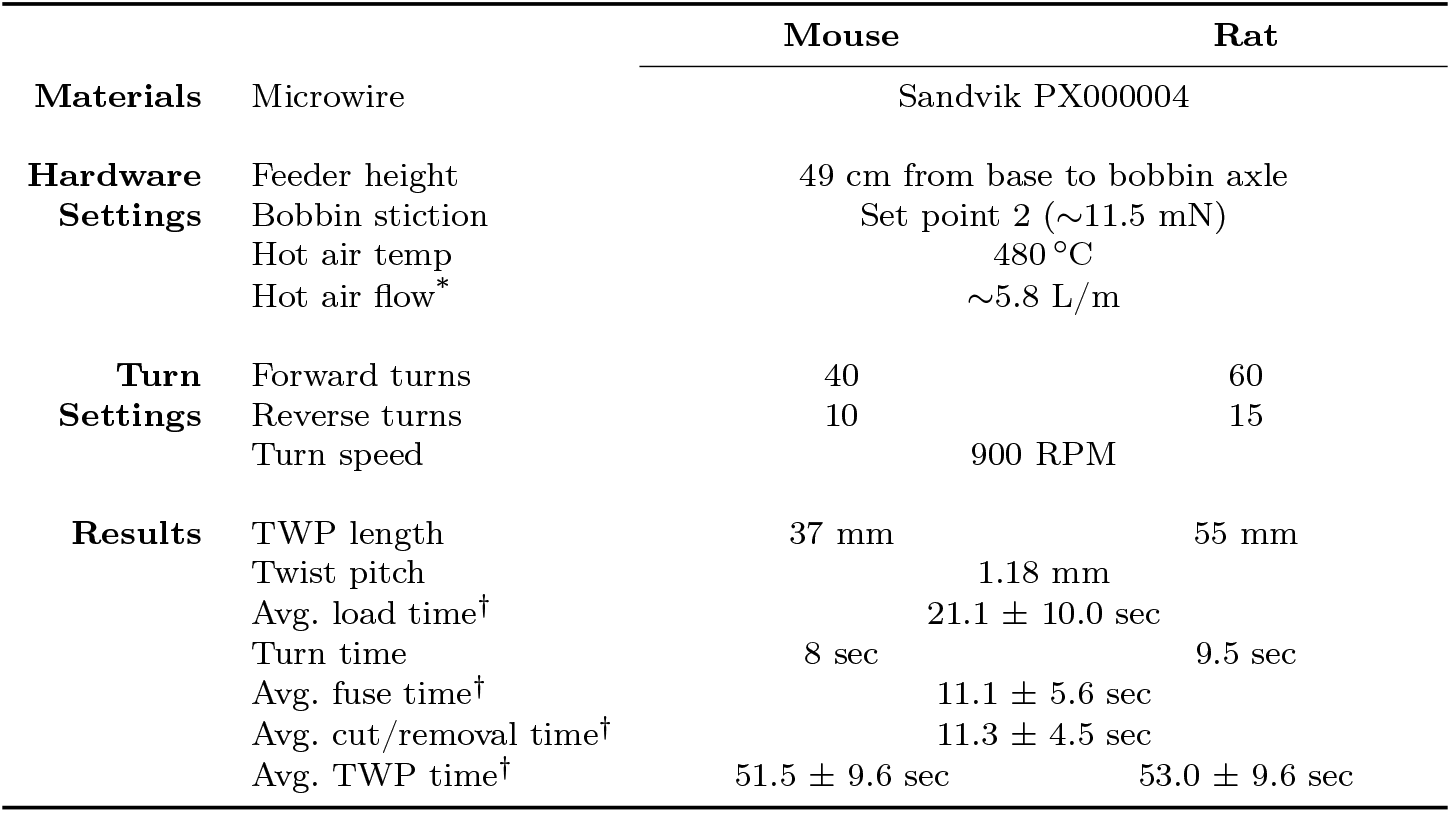
Materials, operation parameters, and resulting tetrode features as Twister3 is used in our labs. Critical settings are the feeder height, which controls the TWP pitch and bobbin torsion, which controls wire tension. * Using the hot air station specified in the bill of materials, this is the lowest setting used in combination with 8 mm diameter nozzle ^†^ Mean ± standard deviation over three novice users (Fig. 5)

Properly loaded bobbins will have microwire tightly wound around their center groove. This relies on carefully adjusting the position of the wire guide such that it’s tip is located about 1–2 mm from the center groove. The loading processes should be monitored as it begins to ensure that microwire is being accepted by the bobbin. If there is an issue, pressing the knob on the control unit will halt the process so it can be corrected.

### 3.3 Making Twisted Wire Probes

The following steps detail tetrode construction using Twister3. If you are interested in making stereotrodes instead of tetrodes, follow the same steps but only use two bobbins on opposing sides of the feeder.

1. Attach the leaf spring to the base rotor (Fig. 3 (B1))
2. Draw the wire down from all of the bobbins and group the tips with your fingers. If one wire is much longer than the others, trim it.
3. Clamp the tip of bundle with the alligator clip. The position of the wire bundle in the clip is not important (Fig. 3 (B2)).
4. Flip the clip 180° such that the wire bundle wraps around the bottom of the clip and exits its rear face (Fig. 3 (B3, inset)). This will ensure that the wire does not slip and that the twist alignment jig will keep the bundle exactly concentric with the axis of rotation during twisting. Keep the clip in your hand while performing the following 2 steps.
5. With your free hand, pull up on the twisting attachment’s leaf spring until under slight tension, about 1.5 cm (Fig. 3 (B3)).
6. Draw the alligator clip down to meet the magnet on the twisting attachment, feeding the bundle into the alignment jig (Fig. 3 (B4)). The alligator clip can be rotated before drawing to wrap the wire around it and improve its grip on the bundle.
7. Ensure all wires are guided through the center of the alignment jig (Fig. 3 (B4, inset)) and the loose ends are not interfering with the taught portion of the wire.
8. Slowly lower the leaf spring until it is in equilibrium with the upward force produced by the tetrode wire. Each of the wires should be pulled straight. If any wire has slack, its bobbin can be turned backwards slightly until it is taut. Do not let the spring snap back under its tension, as this will leave slack in the wires
9. Set the controller to “mode 0”. Set the desired number of turns and turn speed. We use 900 RRM for our wire. This only needs to be done once, or whenever parameter changes are required.
10. Press and hold the knob down to perform a twist.
11. When finished, fuse wires *starting from point at which they separate towards the bobbins* using the hot air gun at ~480°C (Fig. 3 (B5)). We have found that fusing from the bottom will cause the lower portion of the TWP to ‘absorb’ slack from above resulting in a very fine twist pitch and a TWP that is shorter than intended.
12. Using two hands, simultaneously roll each of the bobbins forward a bit in order to release tension on the tetrode wire (Fig. 3 (B6))
13. Cut the tetrode wire above the point at which the wires are fused (Fig. 3 (B7)). Make sure to leave enough free wire for connectorization.
14. Pull the alligator clip off the magnet and cut the finished tetrode into a storage box (Fig. 3 (B8)).

Choosing twisting parameters will require some experimentation in order to produce TWPs with the desired geometric and mechanical properties given the user’s choice of wire, implant type, and animal model. In our labs, we use Twister3 to make tetrodes for microdrive implants in both mice and rats. The operation settings that we use are shown in Table 1. Two settings of note are the height of the wire feeder above the motor assembly, which determines the probe length and microwire twist pitch and the bobbin stiction threshold which determines the wire tension during twisting and drawing. Figure 4 shows the effect of changing the wire feeder height on resulting tetrode characteristics. Although lowering the feeder closer to the motor, and therefore increasing the angle of wire divergence from the axis of rotation does decrease twist pitch and probe length for a given number of turns, the buckling point (Fig. 4) and stiffness (Fig. 4) of tetrodes across feeder heights are remarkably stable.

**Figure 4:**
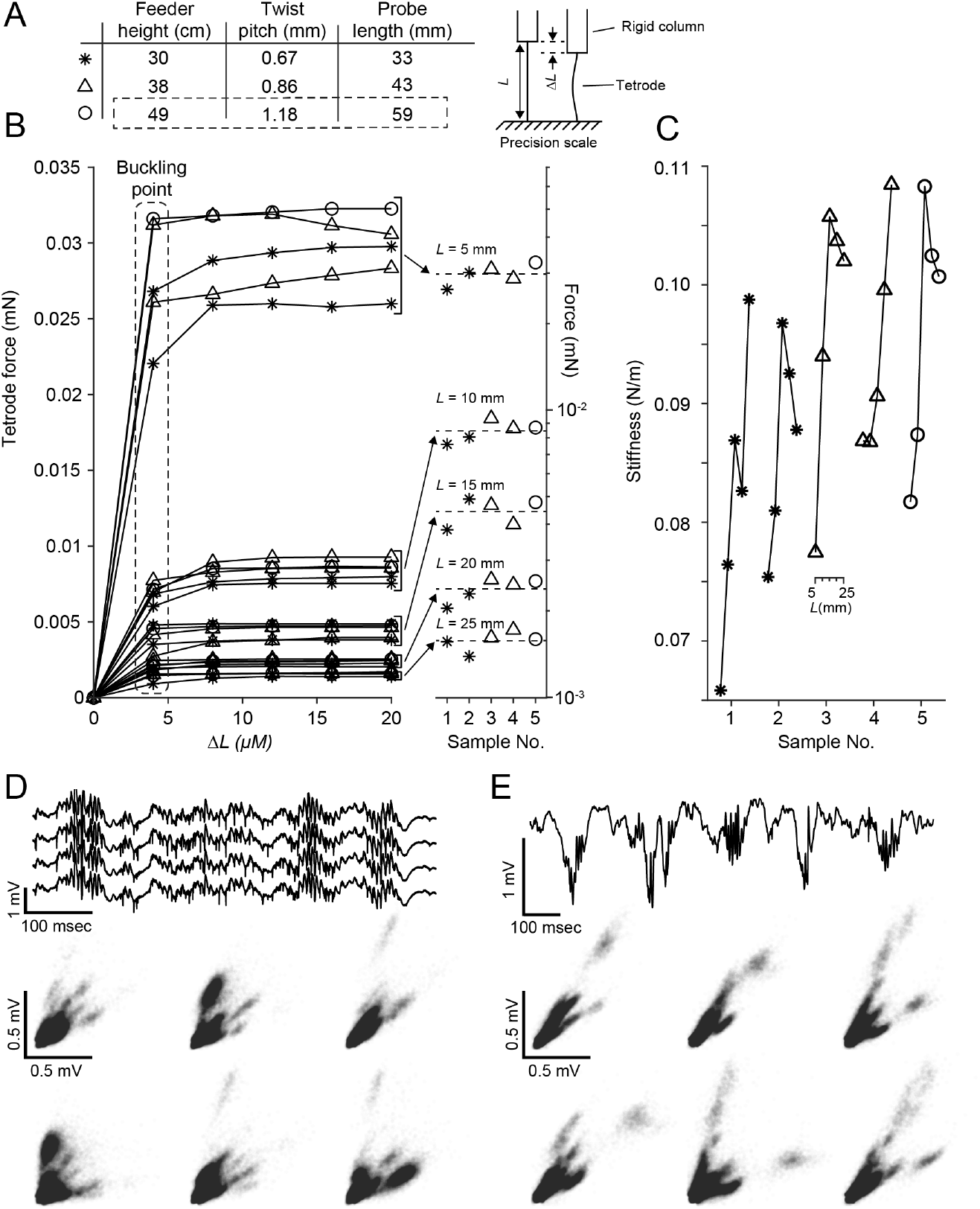
Tetrode characteristics for different twist pitches. **(A)** *(left)* Feeder height versus twist pitch and probe length for the tetrodes in this figure. All other construction parameters were kept the same as Table 1. Circled row indicates the parameters used in the tetrode used to create panel (D). *(right)* Diagram of the mechanical test. Tetrodes were attached to a rigid column and the exposed portion cut to a length *L*. The rigid column was lowered in small increments using a micromanipulator onto a precision scale and the restoring force was measured to find the buckling point. **(B)** *(left)* Compressive force versus depth lowered (*δL*) onto a rigid surface. Each line is a single tetrode sample cut to one of 5 exposed lengths (*L* = 25, 20, 15, 10, and 5 mm). Data point symbols correspond to the table in (A) for various probe lengths. The buckling force (value at which there is no increas in restorative force with *δL*) is length dependent. Different exposed probe lengths form clear groupings with the bucking force increasing as the probe length gets shorter. *(right)* Buckling forces for each sample on a log scale. The buckling force is clearly grouped for each probe length but is not affected by twist pitch. **(C)** Stiffness versus probe length for each of the samples tested. Of the parameters tested, longer, wide-pitch TWPs tended to be stiffest. **(D)** Example hippocampal mouse CA1 recording during sleep. *(top)* Raw voltage trace from a single tetrode. Three sharp wave-ripple events are clearly visible in the trace. *(bottom)* Spike amplitudes for each combination of two wires on the tetrode. **(E)** Example hippocampal rat CA1 recording during sleep. *(top)* Raw voltage trace from a single electrode. Multiple sharp wave-ripple events are clearly visible in the trace. *(bottom)* Spike amplitudes for each combination of two wires on the tetrode. Recordings in (D) and (E) are skull referenced.

To test probe functionality, we produced tetrodes using the device settings in Table 1. Using procedures that were approved by the Committee on Animal Care of Massachusetts Institute of Technology and followed the ethical guidelines of the US National Institutes of Health, these tetrodes were gold plated [12] and used in combination with microdrive assemblies [5, 13] to obtain recordings in the pyramidal cell layer of CA1 of mice and rats. As expected, these tetrodes reliably produced characteristic LFP and multiple, well-isolated units in both mice (Fig. 4(D)) and rats (Fig. 4(E)).

### 3.4 TWP Construction Time

To quantify Twister3’s speed of operation, we measured the tetrode construction time of three users. All users had ~1 hour of experience with the device at the time of testing (Fig. 5). We divided speed measurements into three steps: *(1)* wire clipping/drawing, *(2)* wire twisting/fusing, and *(3)* tetrode removal and storage. Time trials were performed using the device parameters shown the “Mouse” column of Table 1. The motor-in-motion time was constant (9.1 sec at 1000 RPM max speed) and therefore incurred a constant offset on step (2), as indicated by the dashed line in Figure 5. All users were able to create tetrodes at a pace exceeding 1 tetrode/minute. However, each user had slightly different strategies when using the device. For instance, user 3, who has large amounts of experience with tasks requiring fine motor skills, was relatively quick for steps (1) and (3) and was relatively slow for the fusing step (2). User 1 had occasional difficulty clipping and the wire during step (1), increase their average time. These discrepancies indicate that there is room for future improvement and automation, especially with respect to wire clipping and fusing.

**Figure 5:**
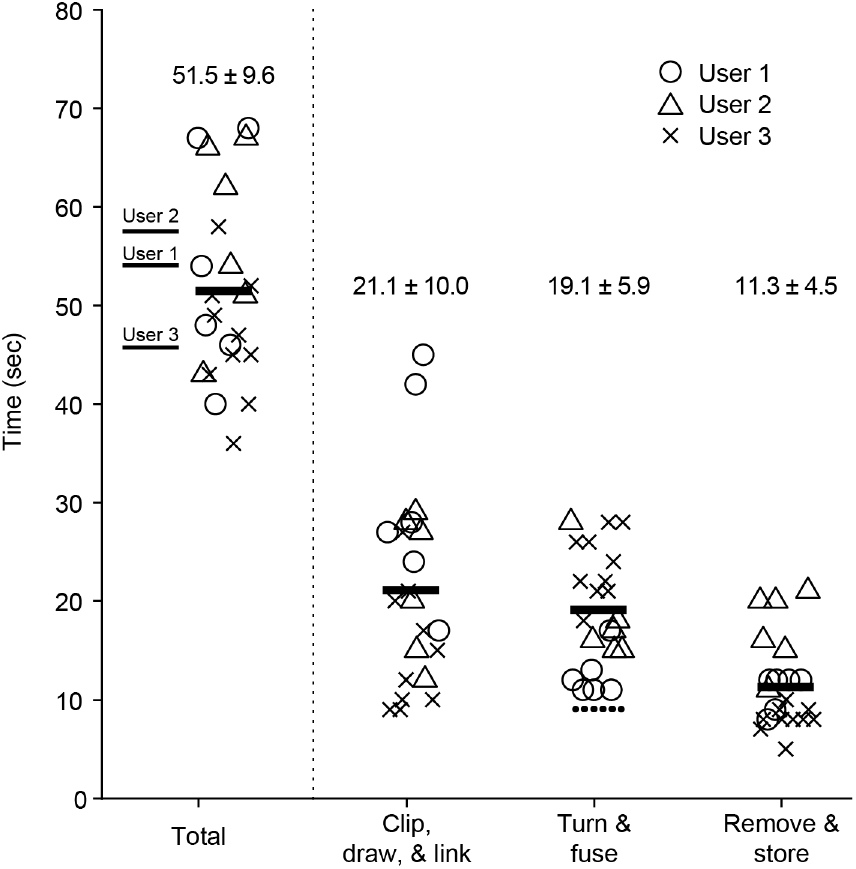
Tetrode construction time of users with ~1 hour of experience with Twister3. *(left)* Total construction time for each tetrode (symbols), average time across tetrodes for each user (thin lines), and across all tetrodes and users (thick line). *(right)* Timing for each step of tetrode construction. The dotted line above the turn and fuse step is the constant motor-in-motion time. Twister3 parameters were the same as the “Mouse” column of Table 1.

## 4 Materials and Assembly

A potentially updated bill of materials (BOM) for electrical, mechanical and 3D-printed parts is available on a Google sheet^11^. Instructions for component assembly are provided in the following sections. For ease of reference and permanence, we provide snapshots of current BOMs in Appendix B. However, the online versions should be used during Twister3 assembly to ensure up to date suppliers and error corrections.

### 4.1 Mechanical Components

The mechanical portion of Twister3 consists of common hardware, standard optomechanical components, and 3D-printed parts. Wherever possible, we used standard (and easy to replace) parts. The mechanical bill of materials is shown in Table 3. In addition to these standard mechanical components, several 3D-printed parts are required (Table 4). These parts are available for direct purchase from third-party 3D-printing services via the links provide in the table. After obtaining the required components, the wire feeder, wire-guide, and stock-spool assemblies can be constructed by following steps detailed Figures 6, 7 (A), 7 (B), and 7 (C), respectively. After each of these modules is complete, they are combined into the complete device by following the steps presented in Figure 8.

**Figure 6:**
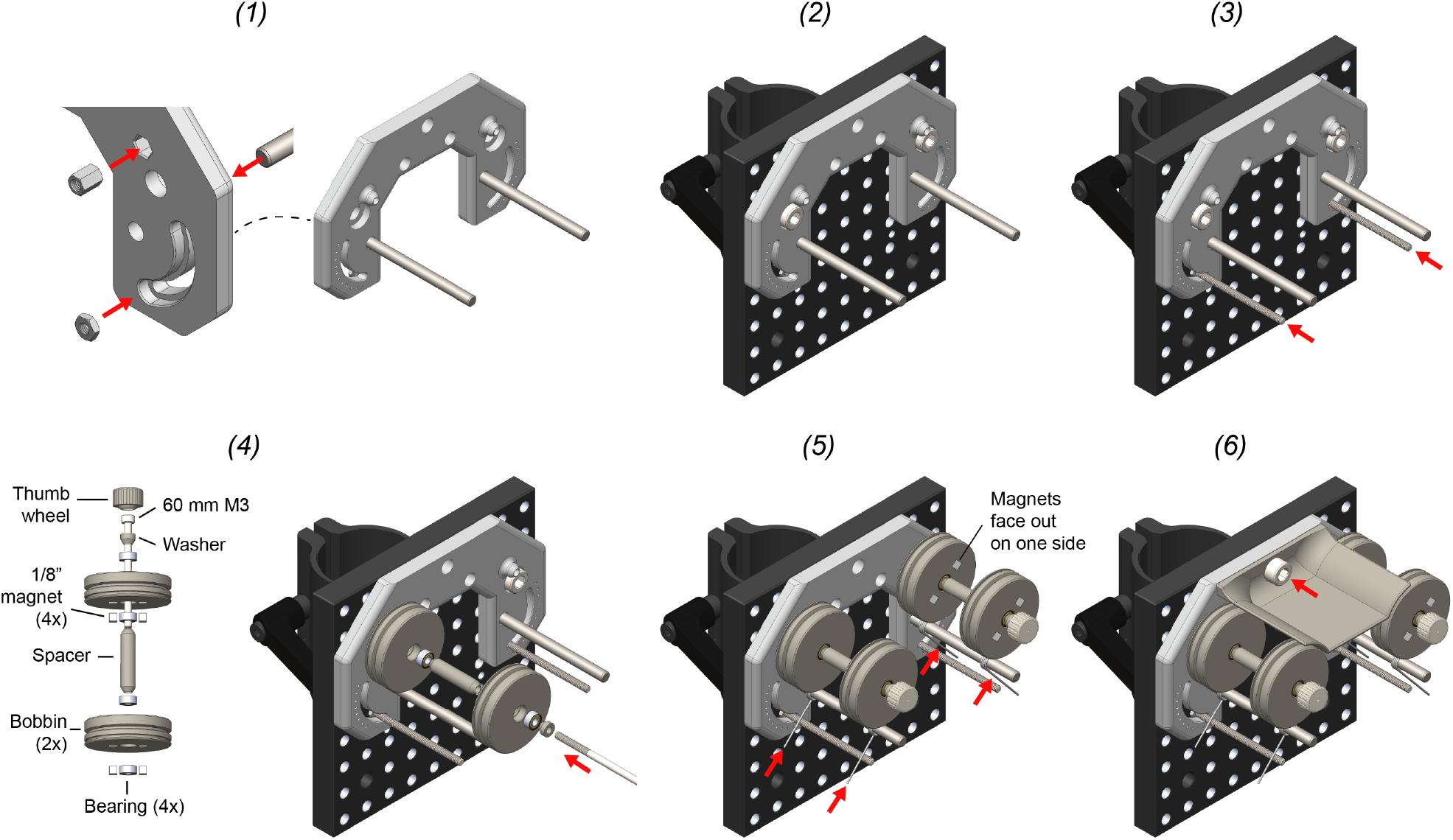
Wire feeder assembly. 1. Insert press-in components into the feeder base. This includes M3 nut (2x), M3 standoff (2x), and 3/16” diameter dowel pin (2x). This requires a mallet. 2. Mount the feeder base unto the C1545/M mounting clamp using M6 screws (2x). The top of the feeder should be flush with the mounting clamp. 3. Cut two 5 cm sections from the M3 threaded rod. Turn each section into the M3 nut which behind the feeder base. The position of the rod determines the stiction on each bobbin during wire draw. Lower positions provide less stiction. We have found that the second notch is a good position to start with. 4. Use the 60 mm M3 screw to mount the bobbin assembly to the standoff captive within the feeder base. Repeat for both sides. The thumb-screw head should be glued onto the M3 screw using epoxy prior to this step. 5. Thread a torsional spring onto the dowel pin. Squeeze it together and then set it between the threaded rod on one side and the shallow groove in each bobbin on the other. Repeat for each bobbin. 6. Install the wire shield above the bobbins using a single M6 screw.

**Figure 7:**
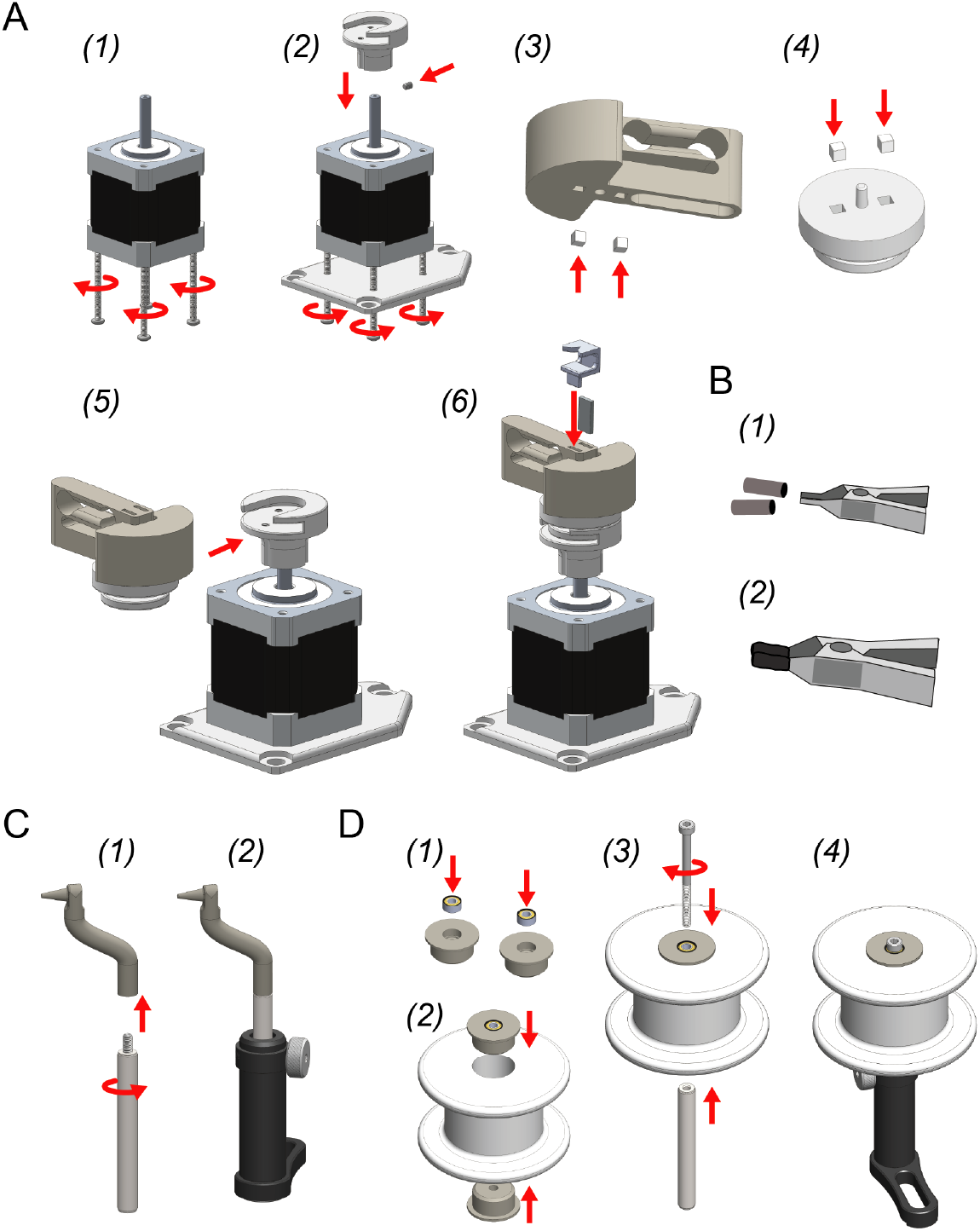
Other mechanical assemblies. **(A) Motor assembly.** 1. Remove the long, M3 step screws from the bottom of the stepper motor (4x). 2. Use 40 mm M3 screws (4x) to attach the motor mount to the bottom of the motor. 3. Fix the rotor base onto the shaft with the M3 set screw. 4. Press fit two magnets into the alignment plate *in the same orientation*. Make sure they are pushed down until recessed below the plastic surface so that they do not interfere with the flat mating surface of the piece. 5. Insert the alignment plate into the slot on the rotor base. 6. Attach two *additional* magnets on top of those you just inserted into alignment plate. Press the spring rotor onto these magnets to press fit them into the spring rotor base. This procedure ensures that magnets will be press fit into the spring rotor base with the correct polarity. Make sure the magnets are recessed below the bottom surface of the spring rotor so that it rests flat on top of the alignment plate. 7. Push the clip magnet into one of the slots on the spring rotor top. In the remaining slot, press the twist alignment jig into position over the magnet. **(B) Wire clip assembly.** 1. Put two pieces of heat shrink tubing over the wire clip jaws. 2. Shrink into position using the hot air gun. This prevents electrode wire from slipping during a draw. The clip can then be stuck under the wire alignment jig (Fig. 3). **(C) Wire guide assembly.** 1. Screw the wire-guide into a mini, 6 mm diameter optical post. 2. Push the optical post into a swivel post holder. **(D) Stock spool assembly.** 1. Push a bearing into each of the stock spool bearing cases. 2. Push each of the bearing cases into the stock spool of tetrode wire. 3. Push a 40 mm, M3 screw through the two bearings and screw into a mini, 6 mm diameter optical post. 4. Push the optical post into a swivel post holder.

**Figure 8:**
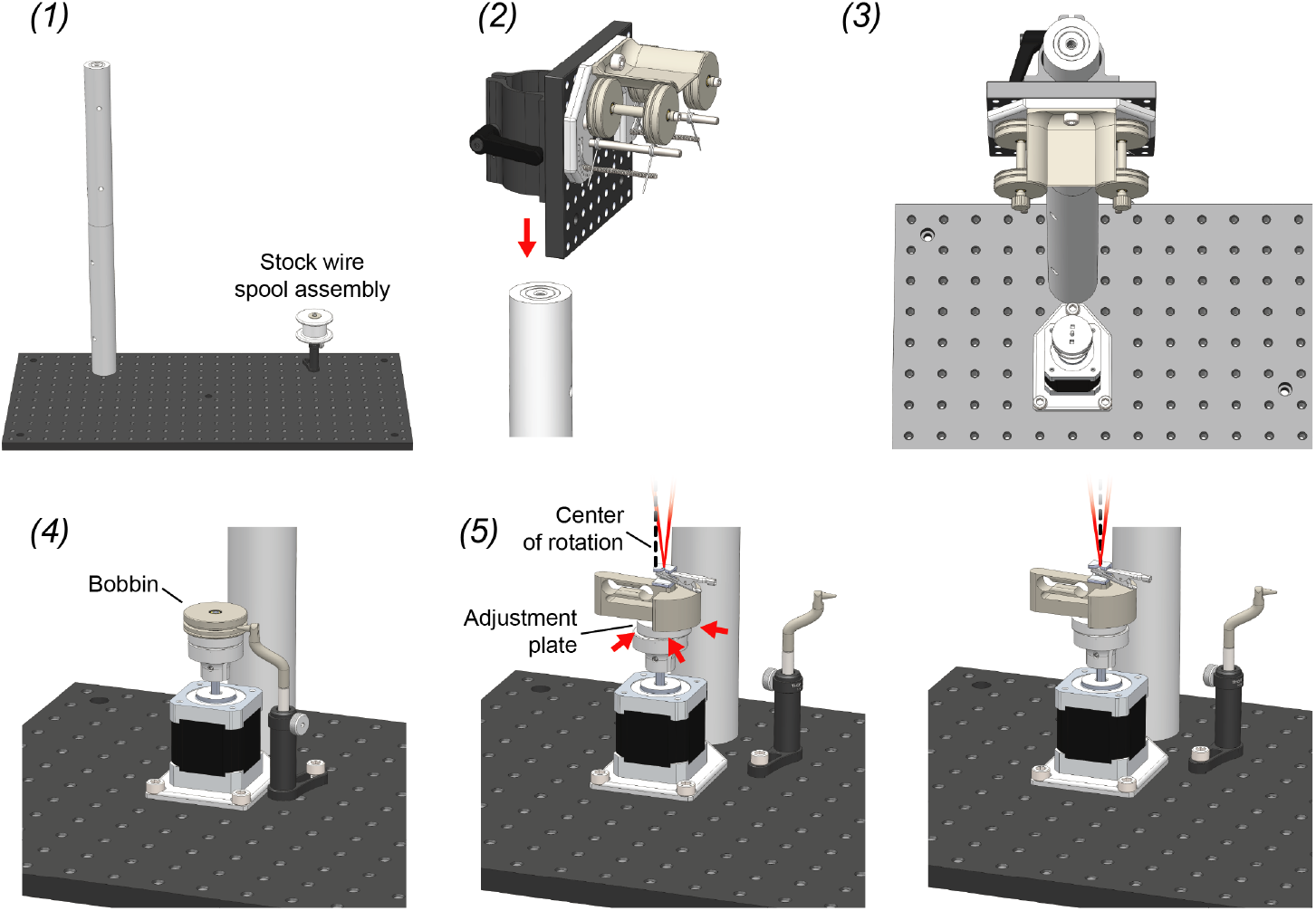
Twister3 assembly. 1. Screw together the large, 1.5” diameter mounting posts and then screw this long post into either the left or right side of the optical bread board. Mount the stock spool assembly in on the opposite side of the optical breadboard using a M6 screw. Its exact position does not matter. 2. Mount the feeder assembly on the post using the post mounting clamp on its back. 3. Mount the rotor assembly directly in front of the post, as close as it will go, using 3 M6 screws. 4. Mount the wire guide assembly into a position that is in close proximity to the motor assembly using a single M6 screw. The tip of the wire guide should be able to extend into the center grove of a wire bobbin when it is mounted on the rotor bases for wire reloading. 5. With the wire-clip mechanism installed, slide the adjustment plate around until the motor axis of rotation (dotted black line) is precisly in line with the wire bundle (red lines). When properly aligned, the apex of the wire bundle will appear motionless during motor turning.

### 4.2 Control Electronics

The control board is comprised of the following blocks: power regulation, motor-driver, microcontroller, and user interface (Fig. 2 (A)). Wherever possible, we used pre-assembled modules (microcontroller, motor driver, and LCD display). The bill of materials for this board is shown in Table 5. Printed circuit board designs and Gerber files are available on the Twister3 repository^12^.

## 5 Discussion

There are several device options for making TWPs. These broadly fall into two categories: *(1)* Twisters with a manual wire folding step, and *(2)* Pre-loaded bobbin designs. Devices such as the Open Ephys Twister^13^, Matt Gaidica’s Simple Twister^14^, and the Neuralynx Spinner-2.0^15^ fall into the first category. The first of these two devices are very cheap and simple, and may be ideal for labs who do not need to make many tetrodes and can accept some TWP construction variability. However, a general disadvantage with manual folding machines is that they are slow. This is due to a manual folding step combined with loose mechanical coupling that requires slow turning speeds. A single TWP generally takes several minutes to make, even for an experienced operator. This can be partially mitigated by using these devices in parallel: a second TWP is folded while the first is turned. However, we have found that due to the finicky nature of the folding step, using any more than two devices at a time is nearly impossible. Further it places a large rote labor burden on the operator, which can lead to poor construction quality due to boredom.

Aside from slow construction speed, these devices introduce large (and uncontrollable) variability in manual wire handling and twist-pitch (which is directly linked to electrode compliance (Fig. 4)). Although they are cheaper than bobbin-based designs, their manual labor requirements and slow operation lead to human-resource requirements that can far outweigh the increased material cost of bobbin-based designs.

There are two options for pre-loaded bobbin TWP machines: Twister3 and the SpikeGadgets Tetrode Machine^16^. Although similar in principle of operation, these two designs use different strategies at nearly every component resulting in very different user experiences and cost. We have summarized these differences in Table 2.

**Table 2:** Comparison of Twister3 and SpikeGadgets Tetrode Machine.

|  | **SpikeGadgets**<br>**Tetrode Machine** | **Twister3**<br>**(this device)** |
| --- | --- | --- |
| **Wire feed mechanism** | Spring-loaded clamp | Torsion-spring quick draw |
| **Automatic wire fusing** | Yes | No |
| **Wire/motor-axis alignment** | No | Yes |
| **TWP turning motor type** | Continuous DC | Stepper |
| **Wire fuser motor type** | Stepper | N/A |
| **Motor controller** | Standard half-bridge | TMC2130 |
| **TWP turning speed** | 60 RPM typical | 700–1000 RPM typical |
| **Acceleration control** | No | Yes |
| **Bobbin/motor attachment** | Screw | Magnets |
| **Wire bundle/motor attachment** | Free-hanging | Leaf-spring & magnet |
| **Microcontroller module** | Arduino Due | Teensy LC |
| **Adjustable twist geometry** | No | Yes |
| **Mechanical parts** | Custom machined | Standard optomechanics & 3D printed |

The two most notable differences between Twister3 and the SpikeGadgets Tetrode Machine are the means by which they increase TWP construction speed and the wire fusing mechanism. Twister3 provides automatic wire bundle alignment and a leaf-spring based bundle to motor coupling, instead of relying on gravity to provide wire tension. This means that wire can be turned at fast speeds while maintaining twist integrity. Because wire turning only takes a few seconds, it is not a rate limiting step in the TWP construction process. Because the SpikeGadgets device lacks Twister3’s motor coupling features, it must turn TWPs relatively slowly (~60 RPM vs. ~1000 RPM).

To compensate for its slow turn rate, the SpikeGadgets machine permits efficient construction via parallelization. Wire is turned slowly, but a linear actuator and hot air gun are used to perform the fusing step automatically. The benefit of this strategy is twofold: (1) increased repeatability of the fusing step, and (2) manual labor is only required for clipping the wire bundle to the motor. This allows 3 identical twisting units to be used in parallel, increasing the effective TWP construction rate to the point where there is effectively no user downtime. The downside of this strategy is a major increase in materials cost and design complexity compared to Twister3. However, given the additional benefits of automated wire fusing, we see this feature as obvious target for future improvement of Twister3.

Aside from these two primary differences, Twister3 also affords several other improvements compared to the SpikeGadgets device. Twister3 provides precise acceleration control, quick-draw wire feeding, rapid magnetic wire bundle to motor attachment mechanism, and automatic wire tensioning. These features simplify operator use in comparison to the SpikeGadgets machine. Further, our use of standard optomechanical and cheap 3D printed parts greatly reduces BOM cost and increases ease of acquisition compared to SpikeGadgets design, which relies on custom, tight-tolerance machined parts.

Because TWPs provide a good balance of data quality, ease of use, ease of assembly, and low cost, they will continue to be used in in vivo neurophysiology labs for years to come. Twister3 is a simple device that greatly decreases manual labor, greatly increases TWP production speed and quality, and is affordable for most labs that might benefit from it. It is fully open-source, well-documented (via this manuscript and instructional videos (Appendix A)), and is composed of easy to obtain parts. Further, we encourage the replication and improvement of this device by others. For instance, an open design that incorporates the automated wire fusing step would further reduce human variability in TWP quality of the current design. We hope that Twister3 complements the growing number of open-source electrophysioloy tools, such as microdrives [13, 5], electrical stimulators [14], optical stimulators [15]^17^, acquisition hardware [16], electrode impedance testers [17], and software [18, 16, 19]. Combined with a growing ecosystem of open-source hardware, e.g. for microscopic imaging [20, 21, 22], DNA amplification [23], audio monitoring [24], culture plate-reading [25], bacterial evolution [26], and closed-loop small animal experimentation [27, 28], these tools will permit labs to be completely outfitted with high-performance, low-cost, open-source tools. We believe this trend will increase the accessibility, transparency, and quality of scientific research in general [29].

## Acknowledgements

The authors gratefully acknowledge Ming-fai Fong for her detailed review of the manuscript. Additionally, we thank Marie-Sophie Helene van der Goes and Dimitra Vardalaki for their helpful feedback on the device design as well as participating in time trial testing. Finally, we thank Hector Penagos for providing the tetrode recordings from mice and rats presented in Fig. 4 of this manuscript.

This work was supported by the Center for Brains, Minds and Machines (CBMM), MIT, funded by NSF STC award CCF-1231216. JPN was supported by the NIH (NRSA 1F32MH107086-01). JV and MTH were supported by funding provided by the NEC Corporation Fund for Research in Computers and Communications at MIT and the NIH (1R01NS106031). JV is a Simons Center for the Social Brain at MIT postdoctoral fellow and MTH is a Klingenstein-Simons Fellow in Neuroscience, a Vallee Foundation Scholar, and a McKnight Scholar.

## Appendix A Instructional Videos

Videos documenting device operation and usage are available on YouTube:

- https://youtu.be/xQbXc738ZuM Annotated video showing how to use Twister3 to load bobbins with electrode wire and how to use Twister3 to make tetrodes.
- https://youtu.be/B0MdM4z-wl0 Video showing the construction of two tetrodes, start to finish.

## Appendix B Bills of Materials

**Table 3:** Mechanical bill of materials. A continuously updated bill of materials is located on this Google sheet.

| *Mechanical Bill of Materials* |  |  |  |
| --- | --- | --- | --- |
| Qty | Description | Supplier | Supplier Part No. |
| 1 | NEMA-17 Stepper Motor | Digikey | 1460-1163-ND |
| 10 | 3 × 7 × 3 mm Bearings | Boca Bearing | MR73-2RSC |
| 1 | M3 x 4mm Set screw | McMaster-Carr | 92605A098 |
| 2 | M3 x 6mm Standoff | McMaster-Carr | 94868A162 |
| 1 | M3 x 0.5 mm Thread Size, 500 mm Long | McMaster-Carr | 90024A219 |
| 5 | M3 x 0.5 mm Thread, 40 mm Long | McMaster-Carr | 91292A024 |
| 2 | M3 x 0.5 mm Thread, 60 mm Long | McMaster-Carr | 92290A131 |
| 8 | M6 x 1 mm Thread, 14 mm Long | McMaster-Carr | 92855A615 |
| 2 | M6 x 1 mm Thread, 25 mm Long, Set screw | McMaster-Carr | 92015A135 |
| 4 | Music-Wire Steel Torsion Spring | McMaster-Carr | 9271K578 |
| 2 | Zinc-Plated, M3 x 0.5 mm Thread Nut | McMaster-Carr | 90695A033 |
| 2 | Dowel Pin 3/16" Diameter, 2-1/2" Long | McMaster-Carr | 98380A520 |
| 2 | ∅1.5" Mounting Post, M6 Taps, L = 250 mm | ThorLabs | P250/M |
| 2 | Swivel-Base Mini-Post Holder, 2" (51 mm) Tall, 1/4" (M6) Slot | ThorLabs | MSL2 |
| 2 | Mini-Series Optical Post, ∅6 mm, L = 50 mm | ThorLabs | MS2R/M |
| 1 | Aluminum Breadboard, 300 mm x 600 mm x 12.7 mm, M6 Taps | ThorLabs | MB3060/M |
| 1 | ∅1.5" Post Mounting Clamp, 112.5 mm x 112.5 mm, Metric | ThorLabs | C1545/M |
| 10 | Neodymium Block Magnets, 1/8" cube | KJ Magnetics | B222G-N52 |
| 2 | Neodymium Block Magnets, 1/2" × 1/4" × 1/16" | KJ Magnetics | B841-N52 |
| 2 | Wire Clip | Digikey | 314-1017-ND |
| NA | Silicone Heat Shrink | Digikey | XX |

**Table 4:**
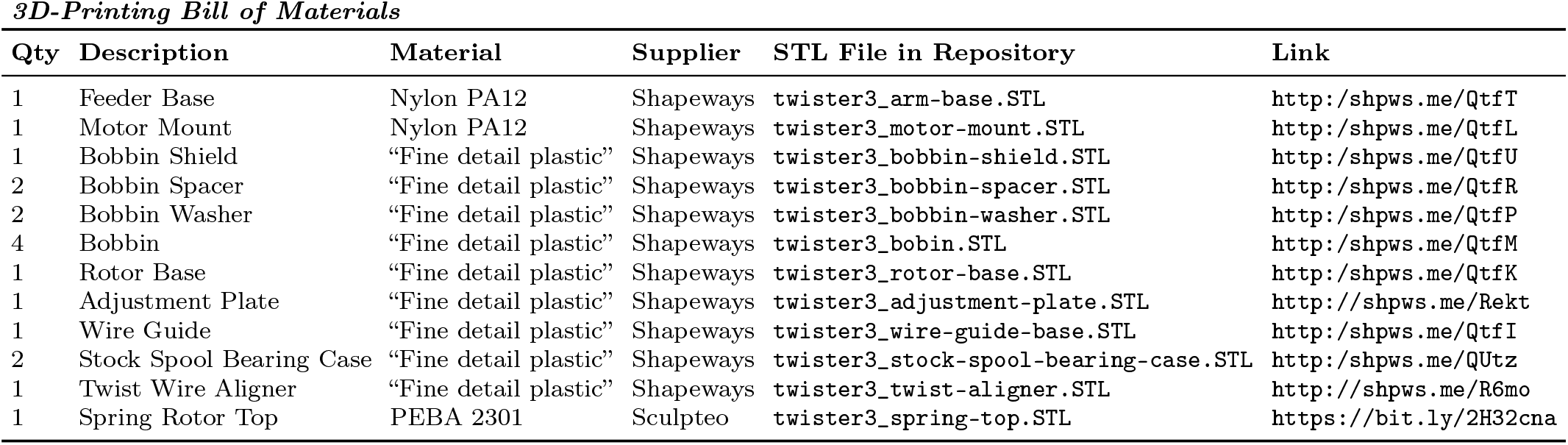
3D-printing bill of materials. An continuously updated bill of materials is located on this Google sheet. The right-most column provides links to order parts from each supplier.

**Table 5:** Control box (Fig. 2) bill of materials. A continuously updated bill of materials is located on this Google sheet. Table entries labeled “common” are ubiquitous and therefore listing specific part numbers is futile due to rapid supplier turn over. PCB design files are located at this link. The PCB is split into two halves separated by a breakaway v-cut. One half serves as the top panel of the enclosure, visible in (Fig. 2 (B)), and the other has all electronics and is mounted beneath the front panel.

| Qty | Schematic Name | Schematic Value | Description | Supplier | Supplier Part No. | Manufacturer Part No. |
| --- | --- | --- | --- | --- | --- | --- |
| 1 | U5 | N/A | Teensy 3.2 | PRJC |  |  |
| 3 | M1, M2, M3 | N/A | M3 Screws, 10 mm | McMaster | 92000A120 |  |
| 1 | N/A | N/A | 24V Wall Adapter | Amazon |  |  |
| 1 | S1 | EC12P | Rotary Encoder | Sparkfun | 10982 | EC12P |
| 1 | S1 | N/A | Knob | Sparkfun | 10597 | P-15x13-T-T2-24A |
| 1 | U6 | GDM1602K | LCD Display 5V | Sparkfun | 709 | GDM1602K |
| 1 | N/A | N/A | Hot-Air Station | Sparkfun | 14557 |  |
| 1 | For U5 | Teensy Header | 2x7 Header | Digikey | 609-3485-1-ND | 61001421121 |
| 2 | For U5 | Teensy Header | 1x14 Header | Digikey | 732-5325-ND | 61301411121 |
| 1 | For U5 | Teensy Socket | 2x7 Socket | Digikey | S7110-ND | PPPC072LFBN-RC |
| 2 | For U5 | Teensy Socket | 1x14 Socket | Digikey | S7012-ND | PPTC141LFBN-RC |
| 1 | For U6 | LCD Header | 1x16 Header | Digikey | 732-5327-ND | 61301611121 |
| 1 | For U6 | LCD Socket | 1x16 Socket | Digikey | S7014-ND | PPTC161LFBN-RC |
| 2 | J10, J11 | Stepper Stick Header | 1x8 Header | Digikey | 732-5321-ND | 61300811121 |
| 2 | J10, J11 | Stepper Stick Socket | 1x8 Socket | Digikey | S7006-ND | PPTC081LFBN-RC |
| 5 | J1, J8, J9 | N/A | 1x4 Socket | Digikey | S7002-ND | PPTC041LFBN-RC |
| 3 | J3, J4, J5 | N/A | 1x4 SMD Header | Digikey | SAM12225-ND | TSM-104-02-L-SV |
| 2 | For U2 | N/A | Board Spacers | Digikey | 952-3465-ND | R6010-00 |
| 3 | M4, M5, M6 | N/A | M3 SMD Nuts | Digikey | 732-10918-1-ND | 9775056360R |
| 1 | J2 | N/A | Screw Terminal | Digikey | ED10563-ND | OSTVN04A150 |
| 1 | J1 | PJ-063BH | Barrel Jack | Digikey | CP-063BH-ND | PJ-063BH |
| 4 | C2, C6, C7, C8 | 10 nF, 50 V | Capacitor, X7R | Digikey | 478-1227-1-ND | 06035C103KAT2A |
| 1 | C16 | 470 nF, 10 V | Capacitor, X7R | Digikey | 1763-1190-1-ND | 0402BW474K160YT |
| 2 | C1, C3 | 1 uF, 50 V | Capacitor, X7R | Digikey | 587-3247-1-ND | UMK107AB7105KA-T |
| 1 | C4 | 15 uF | Tantalum Capacitor | Digikey | 399-3775-1-ND | T491D156K035AT |
| 1 | C5 | 220 uF | Tantalum Capacitor | Digikey | 399-9744-1-ND | T491D227M010AT |
| 1 | R6 | 470 $\Omega$ | Resistor, Thick film | Digikey | Common | Common |
| 2 | R1, R9, R10, R11, R12 | 4.7 kΩ | Resistor, Thick film | Digikey | Common | Common |
| 5 | R2, R3, R4, R5, R17 | 10 kΩ | Resistor, Thick film | Digikey | Common | Common |
| 1 | R13 | 20 kΩ | Resistor, Thick film | Digikey | Common | Common |
| 2 | R8, R16 | 20 kΩ | Trim Pot | Digikey | SM-42TX203CT-ND | SM-42TX203 |
| 1 | L1 | 68 uH | Inductor | Digikey | 513-1386-1-ND | DR125-680-R |
| 1 | D1 | DB2W60400L | Schottky Diode | Digikey | P15184CT-ND | DB2W60400L |
| 1 | U1 | IS31FL3193 | LED Driver | Digikey | 706-1215-1-ND | IS31FL3193-DLS2-TR |
| 1 | U3 | MAX6816 | Debouncer | Digikey | MAX6816EUS+TCT-ND | MAX6816EUS+T |
| 1 | U2 | MIC4680-5.0YM-TR | DC DC Converter | Digikey | 576-1221-1-ND | MIC4680-5.0YM-TR |
| 1 | U4 | TMC2100 | Silent Stepper Stick | Digikey | 1460-1159-ND | N/A |
| 1 | N/A | N/A | Enclosure | Digikey | 36-705-ND | 705 |

## Footnotes

1 e.g. http://neuronexus.com/products/neural-probes/

2 http://www.open-ephys.org/twister

3 https://neuralynx.com/hardware/tetrode-spinner-2.0

4 http://www.spikegadgets.com/main/home.html; Note: M.K. is co-owner and M.B. is an employee.

5 http://www.spikegadgets.com/hardware/tetmachine.html

6 https://github.com/jonnew/twister3

7 https://neuralynx.com/hardware/tetrode-spinner-2.0

8 Polyimide-coated nichrome; ∅12.7*μ*m

9 https://www.pjrc.com/teensy/teensyLC.html

10 https://github.com/luni64/TeensyStep

11 https://bit.ly/2H2a4FD

12 https://github.com/jonnew/twister3/tree/master/control-board

13 http://www.open-ephys.org/twister

14 http://www.open-ephys.org/simple-twister

15 https://neuralynx.com/hardware/tetrode-spinner-2.0

16 http://www.spikegadgets.com/hardware/tetmachine.html

17 https://github.com/jonnew/cyclops

## Notes

https://github.com/jonnew/twister3

http://www.open-ephys.org/twister3

https://docs.google.com/spreadsheets/d/1tdc3wfE6V87q8yqBOQvDj7WylAztkh6_2kuL-YzyB0g/edit#gid=0

https://youtu.be/xQbXc738ZuM

https://youtu.be/B0MdM4z-wl0

